# A resistome survey across hundreds of freshwater bacterial communities reveals the impacts of veterinary and human antibiotics use

**DOI:** 10.1101/2022.07.18.500504

**Authors:** SA Kraemer, N Barbosa da Costa, A Oliva, Y Huot, DA Walsh

## Abstract

Our decreasing ability to fight bacterial infections is a major health concern. It is arising due to the evolution of antimicrobial resistance (AMR) in response to the mis- and overuse of antibiotics in both human and veterinary medicine. Lakes integrate watershed processes and thus may act as receptors and reservoirs of antibiotic resistance genes (ARGs) introduced into the watershed by human activities. The resistome - the diversity of ARGs – under varying anthropogenic watershed pressures has been previously studied either focused on few select genes or few lakes. Here, we link the resistome of ∼350 lakes sampled across Canada to human watershed activity, trophic status, as well as point sources of ARG pollution. A high percentage of the resistance genes detected was either unimpacted by human activity or highly prevalent in pristine lakes, highlighting the role of AMR in microbial ecology in aquatic systems, as well as a pool of genes available for potential horizontal gene transfer to pathogenic species. Nonetheless, watershed agricultural and pasture area significantly impacted the resistome. Moreover, the number of hospitals and the population density in a watershed, the volume of wastewater entering the lake, as well as the fraction of manure applied in the watershed as fertilizer significantly impacted ARG diversity. Together, these findings indicate that lake resistomes are regularly stocked with resistance genes evolved in the context of both veterinary and human antibiotics use and represent reservoirs of ARGs that require further monitoring.

## 1. Introduction

The emergence of antimicrobial resistance (AMR) is a major health challenge of global concern threatening the effectiveness of known antibiotics to treat infectious diseases. Over the past decades we have been facing an antibiotic crisis, as AMR increases faster than the discovery of new antibiotics (Lewis, 2020). As a consequence, a recent systematic analysis estimated that 1.27 million deaths worldwide in 2019 were directly attributable to bacterial AMR in 88 pathogen-drug combinations (Murray et al., 2022). Moreover, the treatment associated with resistant infections results in substantial financial burden to countries’ economy: for example an estimated increase in Canadian healthcare costs of $6 to $8 billion per year by 2050 (Council of Canadian Academics, 2019).

Even though it is a current health concern, AMR and the antibiotic resistance genes (ARGs) causing it are ancient phenomena and readily detected in pristine environments (Martínez, 2008; D’Costa et al., 2011; Bhullar et al., 2012; Perron et al., 2015), where they likely serve complex functions including communication and antagonism in competing and cohabiting bacteria (Goh et al., 2002; Skindersoe et al., 2008). Bacterial AMR results from intrinsic or acquired resistance mechanisms of defense. Intrinsic molecular mechanisms, such as permeability barriers and broad-spectrum efflux pumps, evolved as a general response to nonspecific toxic compounds, and often confer resistance against multiple antibiotics (multidrug resistance) (Surette and Wright, 2017). Alternatively, acquired resistance mechanisms evolved as a targeted response to particular antibiotics, such as compound-specific efflux pumps and enzymes that modify antibiotic molecules (Surette and Wright, 2017). Besides its ancient and environmental origin, the spread and evolution of ARGs in pathogenic and non-pathogenic bacterial populations is directly and indirectly affected by human activities (Martínez, 2008; Gillings and Stokes, 2012).

Antibiotic resistant bacteria (ARBs) and ARGs enter the environment in Canada primarily via two routes. The first is through human-associated antibiotics use both in hospitals, as well as in the general population, causing ARBs and ARGs to accumulate in wastewater, which is collected and treated in wastewater treatment plants (WWTPs) (Reinthaler et al., 2003; Hocquet et al., 2016). Importantly, WWTPs are not specifically equipped to remove ARGs and ARBs, and may in fact constitute hotspots for ARG evolution and spread, due to sub-minimum inhibitory concentrations (MIC) of antibiotics in digestors as well as conditions promoting horizontal gene transfer (Rizzo et al., 2013). While many ARBs and ARGs are reduced in concentration in WWTP effluent compared to the influent, their relative abundance and their genetic context can shift before entry into surface water systems (Reinthaler et al., 2003; Szczepanowski et al., 2009; Rizzo et al., 2013; Hocquet et al., 2016). In addition, biosolids from WWTPs have been increasingly used as agricultural fertilizers due to their high nutrient content such as phosphorous and nitrogen (Lu et al., 2012). Biosolids are enriched with ARGs and runoff from biosolid-fertilized fields can thus introduce those genes into neighbouring environments (McLain et al., 2017).

The second route of ARG introduction into the environment is via agricultural and specifically animal feeding operations (AFOs). Despite recent attempts to curb the use of antibiotics in agricultural settings in many countries, for example via bans on feeding antibiotics as growth promotors in Europe and Canada (Kraemer et al., 2019), vastly more antibiotics are used in veterinary than in a human medicine settings; 80% of antibiotics sold in the US and 78% of those sold in Canada are for animals (Ventola, 2015; Public Health Agency of Canada, 2022). ARBs, ARGs, as well as excreted antibiotics are present in animal manure and urine, which is subsequently degraded and used as fertilizer. While manure fermentation decreases the abundance of some ARBs and ARGs, ARGs associated with veterinary antibiotics use can be readily detected in soil after fertilization (Lee and Lee, 2011; Graham et al., 2016).

Large scale human land use such as mining and agriculture can also indirectly increase the presence of ARGs in natural bacterial populations. Contamination with heavy metals, biocides and pesticides may increase ARG frequency in bacterial communities due to co- or cross-resistance of genes involved in the defense mechanism against these contaminants and antibiotics (Baker-Austin et al., 2006; Pal et al., 2015; Yang et al., 2017; Liao et al., 2021).

Although *a priori* AMR of environmental, non-pathogenic bacteria is not a concern to human health, it increases the risk of spreading ARGs to pathogens through horizontal gene transfer (Gillings and Stokes, 2012). For this reason, the surveillance of natural environments is one of the goals of the World Health Organization global action plan on AMR (World Health Organisation, 2017) as it contributes to the assessment of AMR dissemination risk (Berendonk et al., 2015). Freshwater environments in particular may act as ARGs reservoirs (Zhang et al., 2009), as surface waters receive runoff or effluent from important AMR hotspots, such as wastewater treatment plans and manure-fertilized fields (Rizzo et al., 2013; Kraemer et al., 2019).

Therefore, contamination of freshwater system with ARGs and the antibiotic resistant bacteria carrying them has received increased interest. For example, ARGs of interest have been tracked along rivers and in lakes receiving point sources of pollution (e.g., WWTPs or AFOs) (Czekalski et al., 2014, 2015; Storteboom et al., 2010; Pruden et al., 2012; Thevenon et al., 2012; Chu et al., 2018). The absolute abundance of ARGs (and specifically *sul*1 (Pruden et al., 2012; Czekalski et al., 2015)) often decreased as distance from a potential pollution source increased. In addition to such specific point sources of pollution, lakes may receive ARGs and ARBs due to general human activity in the watershed. For example, higher concentrations of all or specific ARGs may be found in lakes with highly urbanized watershed (Yang et al., 2017) due to stormwater runoff containing ARGs (Białasek and Miłobędzka, 2020). Similarly, runoff from fields fertilized with manure or pastures could have similar effects in highly agricultural watersheds (Chee-Sanford et al., 2009). Anthropic activity is often related to the trophic status of a lake. Recent work has linked nutrient status to lake resistomes (Huang et al., 2019; Pan et al., 2020; Rajasekar et al., 2022), including studies showing a correlation between increasing eutrophication and ARG abundance (Thevenon et al., 2012; Wang et al., 2020).

Most studies on natural resistomes in lakes have been limited to one (Chen et al., 2019; Stange et al., 2019; Wang et al., 2020) or few (Pal et al., 2015; Wang et al., 2020) lakes (but see Czekalski et al. (2015)). Moreover, these lakes are often highly eutrophic and impacted by multiple potential ARG pollution sources at the same time, making generalization difficult. Here, we conducted a comprehensive profiling of relative ARGs abundance and diversity in bacterial communities of approximately 350 lakes under varying pressure by human activities across Canada using metagenomic high throughput sequencing data. Lakes are the source of approximately 90% of drinking water in Canada (Environment Canada, 2011) and occupy 10% of the country’s territory (Huot et al., 2019). We sought to understand how human impact affects the presence of ARGs in these environments. We found that, even though there was a vast diversity of natural ARGs independent of human impact, agriculture, and specifically manure applied as fertilizer in a watershed, as well as effluent from hospitals and WWTPs had a measurable input on lake resistomes across Canada.

## 2. Methods

### 2.1 Sampling, DNA extraction and sequencing

We sampled 366 lakes across twelve Canadian ecozones during the summer of 2017-2019 at the height of summer stratification in the context of the NSERC Canadian Lake Pulse Network (referred to as LakePulse below) project as described previously (Huot et al., 2019; Kraemer et al., 2020). Briefly, an integrated epilimnion sample was collected at the deepest point of the lake as determined using a depth sounder and stored in a cooler until further processing on the same day. Subsequently, approximately 500 mL of water were filtered on site onto a 0.22 µm filter after pre-filtration through a 100 µm mesh. Filters were immediately frozen at -80 °C and transferred to the lab. DNA was extracted using the DNAeasy PowerWater kit (QIAGEN, USA) (see Kraemer et al. (2020) for details). DNA was visually inspected for intactness using agarose gels and quantified using a Qubit assay (Lumiprobe, USA).

Extracted DNA was sequenced using the NovaSeq Illumina technology with an insert size of 250 base pairs and read lengths of 150 base pairs at Genome Quebec. Sequenced reads were demultiplexed and trimmed to remove low quality bases and adapters using Trimmomatic v.0.38 with default settings (Bolger et al., 2014). Datasets are available under PRJEB29238.

### 2.2 Environmental parameters

The investigated lakes are a subset of approximately 660 lakes which were analyzed in the context of the LakePulse project (Huot et al., 2019). Briefly, lakes were selected for sampling across twelve Canadian ecozones with lake size and average human impact index (HII) within each ecozone as stratifying factors. Lakes further than 1 km from the closest road were excluded. To calculate a synthetic Human Impact Index (HII) variable, the watershed of each lake was delineated based on GIS data and land use classes assigned to each pixel within the watershed. Land use classes included natural landscapes, pasture, forestry (recent clear-cuts), agriculture, urban (buildings and roads) and mines. To calculate HII, land use classes were weighted differentially (urban, agriculture, mines: 1; pasture, forestry: 0.5; natural landscapes: 0) for each lake.

Population density (pop_wshd) in the watershed was based on WorldPop (https://www.worldpop.org/doi/10.5258/SOTON/WP00645, accessed on 10.03.2022), taking into account the respective year of sampling (2017, 2018 or 2019). We summed all the pixel values (population counts) falling inside each lake watershed to approximate population counts for each lake. For shared watersheds between United States and Canada, we used WorldPop population data from both territories. The number of hospitals (nbr_hospitals) and the number of hospital beds (nbr_beds) in a watershed were based on a hospital census provided by the Public Health Agency of Canada (Public Health Agency of Canada, 2015).

Reported WWTPs (nbr_wwtp) was based on the Government of Canada’s list of Wastewater Systems Effluent Regulations as of the 20th of June 2022 (https://open.canada.ca/data/en/dataset/9e11e114-ef0d-4814-8d93-24af23716489). The average effluent volume per day (in m3) was normalized to the total lake volume in million cubic meters (effluent_per_volume). WWTPs and hospital locations falling in a 1 km buffer area around the watershed boundaries were visually inspected. For those points, we corrected GPS locations when necessary to make sure that they fell on the correct side of the boundary. After GPS corrections, we excluded the hospitals and WWTPs outside the watershed areas. When falling within several watershed areas, we attributed the closest lake to each WWTPs and hospitals.

Animal density (anim_den) and the application of manure-based fertilizer onto fields within a watershed were calculated based on the 2016 Agriculture and Pasture Census published by Agriculture Canada (https://open.canada.ca/data/dataset/ae77c7bc-5289-47fe-ad48-5edd1d11f3d5). Total_manure represent the theoretical fraction of the watershed area where any type of manure was applied in the year prior to the census (Oliva et al., 2022). The mean density of animals in the watershed was calculated as follow: (1) we extracted the agriculture and pasture pixels from the census areas and we redistributed the animal census counts per pixel; (2) we summed the pixel values (*i*.*e*., animal unit counts) falling inside each lake watershed to approximate the total number of animal units per watershed; (3) the total number of animal units was divided by the watershed area. All types of livestock were taken into account and each type of animal was normalised to animal units using a formula based on Beaulieu and Bédard (2003), to ensures that animals with different environmental impact are weighted appropriately.

The trophic state of each lake was defined based on the concentration of TP based on the thresholds for Canadian freshwater systems as follows: ultraoligotrophic (TP<4 µg/L), oligotrophic (4-10 µg/L), mesotrophic (10-20 µg/L), mesoeutrophic (20-35 µg/L), eutrophic (35-100 µg/L) and hypereutrophic (>100 µg/L) (Garner et al., 2022). We removed 17 hyper-saline lakes from the analysis following Garner et al. (2022), leaving a dataset of 349 lake.

### 2.3 Screening for antibiotic resistance genes in unassembled reads

To screen unassembled reads for the presence of resistance genes, all trimmed forward reads were blasted against a translated version of the CARD database as implemented in the args-oap software (Yin et al., 2018) using diamond blast with a minimum id of 80 and a minimum hit length of 25 amino acids (Buchfink et al., 2015). Only the best hit for each read was subsequently analyzed. To estimate the number of cells per sample, we also blasted all forward reads against a database of 30 single copy marker genes as implemented in args-oap (Yin et al. 2018). We calculated the total coverage of each marker gene by summing up the length of the blast hits and normalized this by dividing by the total length of the marker gene and then calculated the estimated cell number of each sample as the average coverage across the 30 marker genes. The normalized relative abundance of an ARG was calculated as the total hits, divided by the length of the gene and the estimated cell number in a metagenomic sample.

### 2.4 Screening for antibiotic resistance genes in assembled metagenomes

Trimmed reads for each lake were individually assembled using the spades assembler v. 3.13.0 with kmers 21, 33, 55, 77, 99, 127 and the assembler_only option (Bankevich et al., 2012). Subsequently, contigs were screened for the presence of ARGs using abricate v.1.0.1 against the CARD database (Seemann, 2020). The output was filtered for a minimum gene coverage of 70 and all scaffolds smaller than 1 kb excluded from further analysis. We investigated the genomic context of the remaining scaffolds using plasClass with a probability cut-off of 50% (Pellow et al., 2020). The taxonomy of ARG-containing scaffolds was assigned using Kraken 2 (Wood et al., 2019).

### 2.5 Statistical analyses

We utilized base R v.4.1.2 to test for interactions between the total normalized relative abundance of ARGs and land use and source variables (R Development Core Team, 2009). P-values were corrected for multiple testing using the Hochberg method. The R packages vegan (Oksanen et al., 2016) and ape (Paradis and Schliep, 2019) were used for analyses of beta-diversity and ggplot for visualizations (Wilkinson, 2011).

## 3. Results

### 3.1 Lake physicochemical parameters

In this study, we focused on 349 Canadian lakes which were sampled in the context of a large survey of lakes across Canada (Huot et al., 2019) (Figure 1). Lakes were sampled between Canada’s three coasts (43-68 °N, 53-141 °W) and represent a broad range of lake surface areas (0.005-100 km^2^) and maximum depths (0.7-150 m). They are located within 12 Canadian ecozones, which are terrestrial regions defined by characteristic environmental conditions following the Canadian Council on Ecological Areas (2014). Following their diverse geographical and ecological settings, the chosen lakes have a wide variety of human activity within their watersheds (0-98% of the watershed impacted by human activity) and range from hyperoligotrophic to hypereutrophic. We identified 32 WWTPs and 21 hospitals in 17 and 12 watersheds, respectively. Details on environmental parameters for all lakes in this study are available in Supplemental Table 1 and at https://lakepulse.ca/lakeportal/.

**Figure 1:**
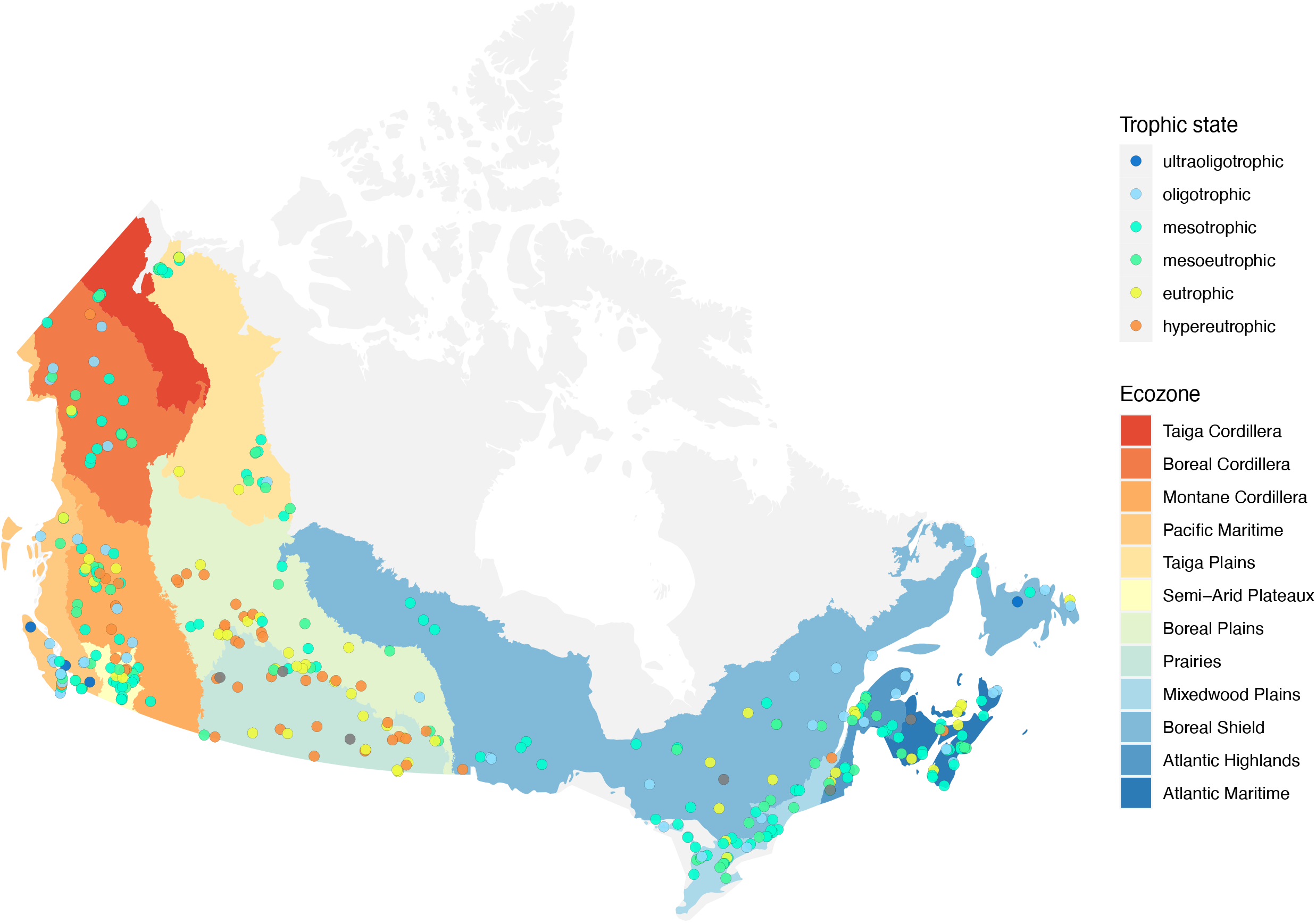
Map showing the locations of the 349 lakes in this study. The trophic state of each lake at the time of the sampling is indicated by the colour of the point.

### 3.2 Antibiotic resistance genes in unassembled read data

Unassembled metagenomic read data offer the opportunity to study the relative abundance and distribution of resistance genes in our samples. Individual lake metagenomes consisted of 17,500K to 91,200K paired reads. We screened the resistomes of 349 lakes by matching them against the CARD database. In total, we detected 468 ARGs, belonging to 20 different resistance gene classes. The ARG class with the highest average relative normalized abundance conferred resistance against bacitracin (average normalized abundance: 7.7×10^−2^), followed by multidrug (3×10^−2^) and Macrolide-Lincosamide-Streptogramin (MLS) resistance genes (4.1×10^−3^). We also detected resistance genes against synthetic antibiotics including sulfonamides, quinolones and trimethoprim, which had lower average relative normalized abundances between 1×10^−5^ and 1×10^−4^ (Supplemental Table 2, Supplemental Figure 1).

#### 3.2.1 Land use impact on the resistome

Across the scales investigated in this study, anthropogenic impacts within the watershed are complex and can originate from a number of factors including: agricultural activity such as fertilisation with manure; urban stormwater runoff; and mining. To normalize these complex pressures on a continental scale, we utilized a synthetic variable called human impact index, which summarises the relative proportion of different watershed land use. Subsequently, we defined a human impact index between 0 and 0.1 as low, between 0.1 and 0.5 as moderate and above 0.5 as high (Kraemer et al., 2020). Overall, the total normalized relative abundance of resistance genes in lake metagenomes was highest in lakes with the smallest anthropogenic impact in their watershed (average normalized abundance 0.134), followed by moderately (0.119) and highly (0.1) impacted lake metagenomes (Kruskal-Wallis test, *p* < 0.001). This result was heavily driven by the highly abundant resistance gene classes bacitracin and multidrug ARGs, which decreased from low to highly impacted lakes (Figure 2). However, for some individual resistance gene classes we observed an increase of the normalized relative abundance. This was the case for aminoglycosides, puromycin, vancomycin, as well as resistance against the synthetic antibiotic trimethoprim (Figure 2).

**Figure 2:**
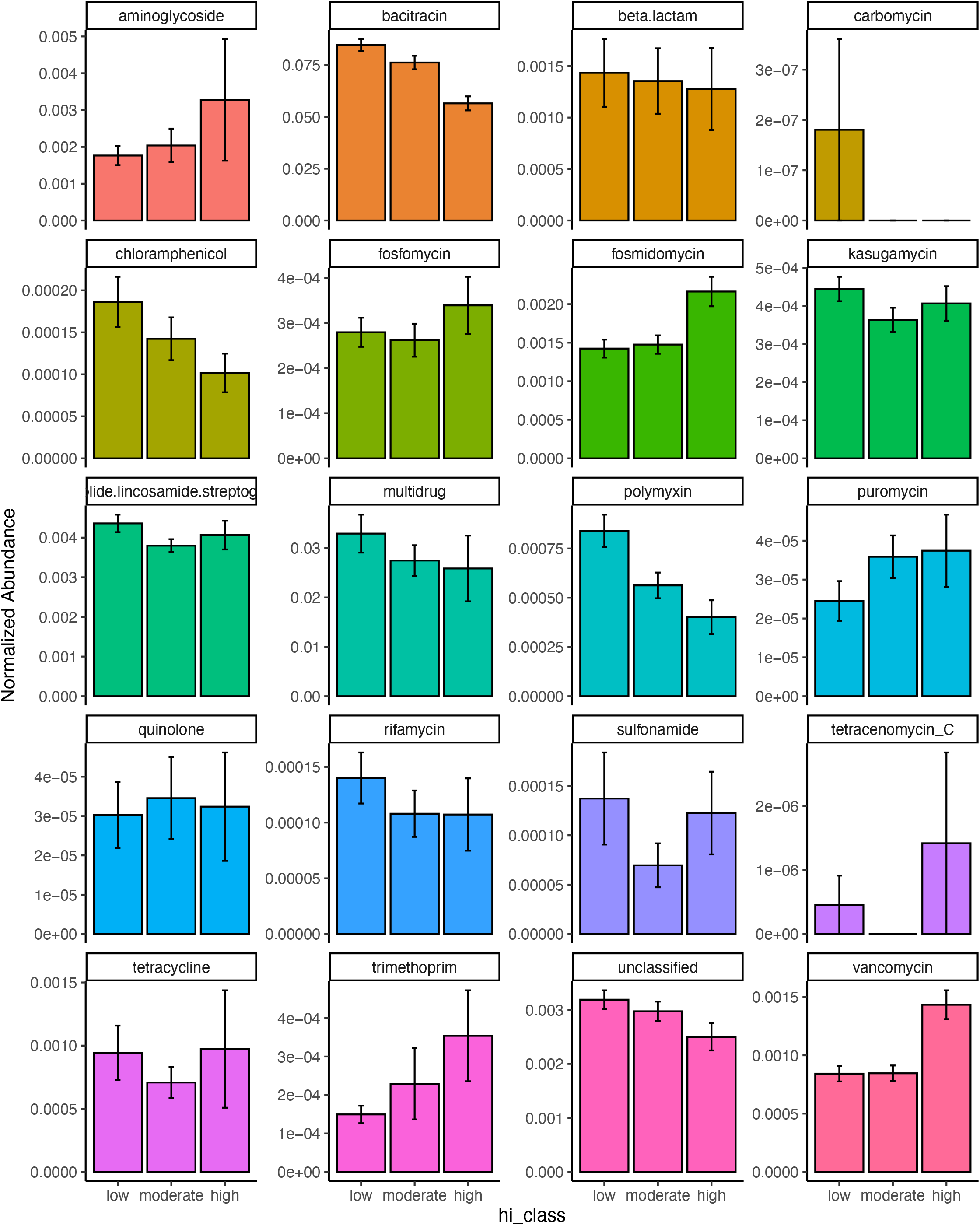
Read abundances normalized by gene length and estimated number of genomes per sample for 20 different classes of ARGs grouped by samples from lakes with low (HII: 0-0.1), moderate (HII: 0.1-0.5) or high (HII: 0.5-1) anthropogenic watershed impact. Error bars represent 95% confidence intervals.

Next, we investigated how HII impacted ARG beta-diversity. We detected a significant impact of HII, as well as lake area and ecozone on the diversity of resistance genes (Permanova, all *p* < 0.05). We explored the structure of lake resistomes further using PCoA. Environmental variables that significantly covaried with the resistome structure were the percentage of agriculture and pasture within the watershed, which contrasted with lakes whose watersheds were characterized by high percentages of forestry (recent clear-cuts) and natural landscapes (Figure 3A). In contrast, we did not observe a significant structuring of ARG classes with land use. In fact, the vast diversity of beta-lactam and multidrug efflux pump resistance genes was largely independently distributed from any of the land use variables measured (Figure 3B). The sulfonamide resistance genes *sul*1 and *sul*2 have been previously investigated as markers of anthropogenic pollution (Czekalski et al.; Pruden et al., 2012). Here, we found that these genes covaried with the second PCoA axis but were not correlated directly with agriculture or pasture impact on the lake resistome (Figure 3B).

**Figure 3.**
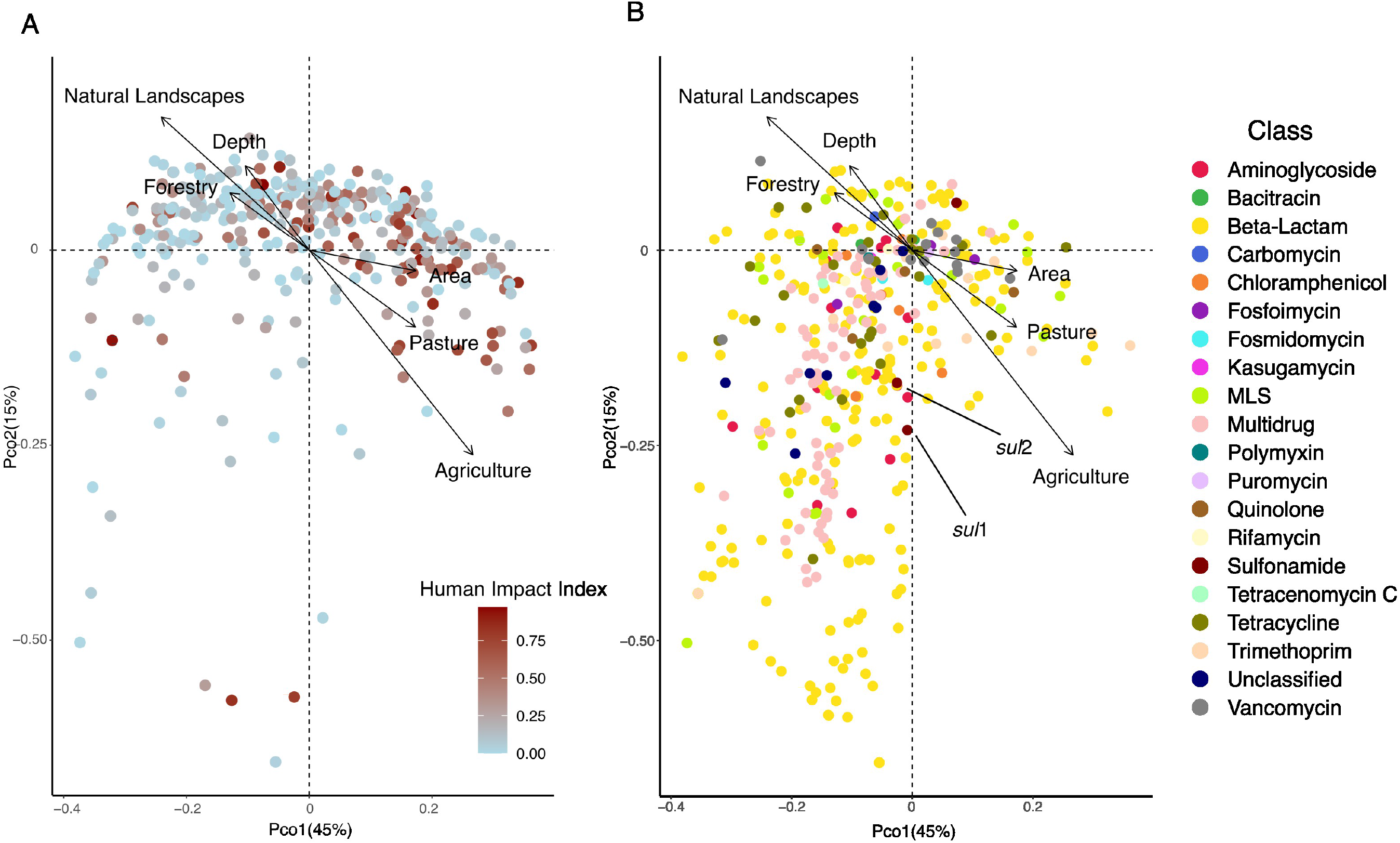
A) PCoA showing the distribution of 349 lake samples projected in the first two dimensions. Each sample is coloured by its human impact index (HII, see text). Arrows show variables that significantly correlated with the ordination after environmental fitting (tested variables: fraction agriculture, pasture, mining, urban, forestry, natural landscapes in the watershed). B) PCoA showing the distribution of individual resistance genes (N=468) projected in the first two dimensions. Each resistance gene is coloured by its class. Arrows show variables that significantly correlated with the ordination after environmental fitting.

#### 3.2.2 Trophic state impact on the resistome

The HII variable represents a relatively static description of the potential state of a lake. A more direct way to assess lake state is to assign trophic status based on the available total phosphorous measured *in situ* at the time of sampling. We were able to assign a trophic status to 343 out of the 349 lakes. Four lakes were ultraoligotrophic, 46 oligotrophic, 122 mesotrophic, 67 mesoeutrophic, 58 eutrophic and 46 hypereutrophic at the time of sampling. In accordance with the HII impact on relative ARG abundance, we found that trophic status correlated with the total normalized abundance of resistance genes, such that oligotrophic (and even the under-sampled ultraoligotrophic) lakes had a higher normalized abundance which steadily decreased up to hypereutrophic lakes (Kruskal-Wallis test, *p* < 0.001). This pattern was driven by bacitracin ARGs (Figure 4). Other resistance gene families showed varying patterns in correlation with lake trophic status. For example, beta-lactam, quinolone and tetracycline ARGs appear to peak in mesotrophic lakes, whereas trimethoprim, vancomycin, fosmidomycin and puromycin increase with increasing lake trophic status (Figure 4).

**Figure 4:**
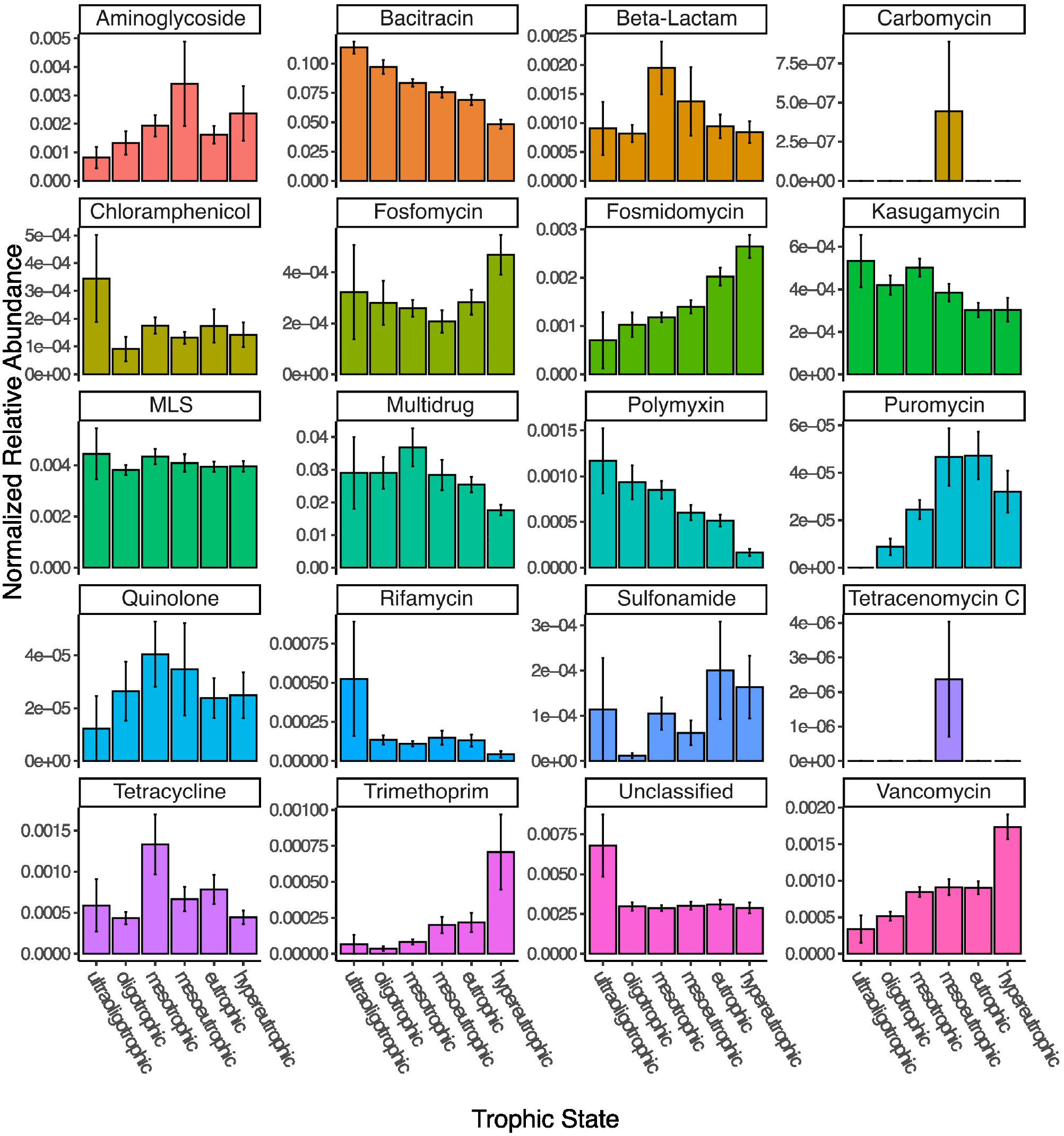
Normalized read abundances (by gene length and estimated number of genomes per sample) for 20 different classes of ARGs grouped by samples from lakes with different trophic status.

While lake trophic status significantly correlated with lake resistome beta diversity (Permanova, *p* < 0.001), it explained only approximately 10% of between-lake variation (distance-based redundancy analysis, *p* = 0.001).

#### 3.2.3 Potential sources of resistance genes

Resistance genes in natural environments can be either autochthonous, or allochthonous and derived from sources of pollution such as animal production or human antibiotics usage. To investigate whether we could detect the impact of distinct pollution sources on lake resistomes, we considered two variables related to veterinary antibiotics use and five variables related to human influence. Veterinary use-related variables are the density of farm animals per census agriculture area as well as the fraction of the watershed to which manure was applied. Variables pertaining to human antibiotics usage are the number of hospitals and hospital beds, the number of WWTPs, the wastewater effluent normalized to lake volume, as well as the total mean density of the population within the watershed. We did detect a significant impact of manure applied in the watershed onto ARG beta-diversity (Permanova, all *p* > 0.05), but did not detect any significant correlations between total normalized abundance and the source variables tested.

Next, we investigated whether the source variables impacted resistance gene diversity by converting the normalized relative abundances to presence-absence data. Watershed population density, number of hospitals, wastewater effluent per lake volume, as well as the total watershed fraction of manure applied were all found to significantly structure resistome diversity (Permanova, all *p* < 0.05). Moreover, we detected significant positive correlations between the number of hospitals and hospital beds within a watershed and the number of different resistance genes detected in a lake metagenome (Pearson’s product-moment correlation, cor = 0.21 and 0.23, all *p* < 0.001), as well as the normalized effluent volume (cor = 0.16, *p* = 0.015), while the number of WWTPs was marginally correlated (cor=0.13, p=0.08). In total, sources of pollution explained only 5% of variation in ARG diversity (Permanova: unexplained variation 95%). Fitting of environmental variables after PCoA revealed a significant fit to the resistome diversity for the number of hospitals (*p* = 0.003), hospital beds (*p* = 0.001), animal density (*p* = 0.04), and manure used in the watershed (*p* = 0.001) as well as a marginal fit for the volume of wastewater effluent (*p* = 0.06) and the number of WWTPs (*p* = 0.98) (Figure 5A-B). Human antibiotics use-associated variables were oriented perpendicularly to the fit for the density of animals in the watershed, indicating how these different sources of pollution influence the resistome in distinct ways. Moreover, fitted arrows for the number of hospitals/ hospital beds and the volume of wastewater are associated with the first PCoA axis and are associated with many of the beta-lactam and multidrug efflux pump genes (Figure 5), which were previously not linked with any of the land use variables tested (Figure 3).

**Figure 5:**
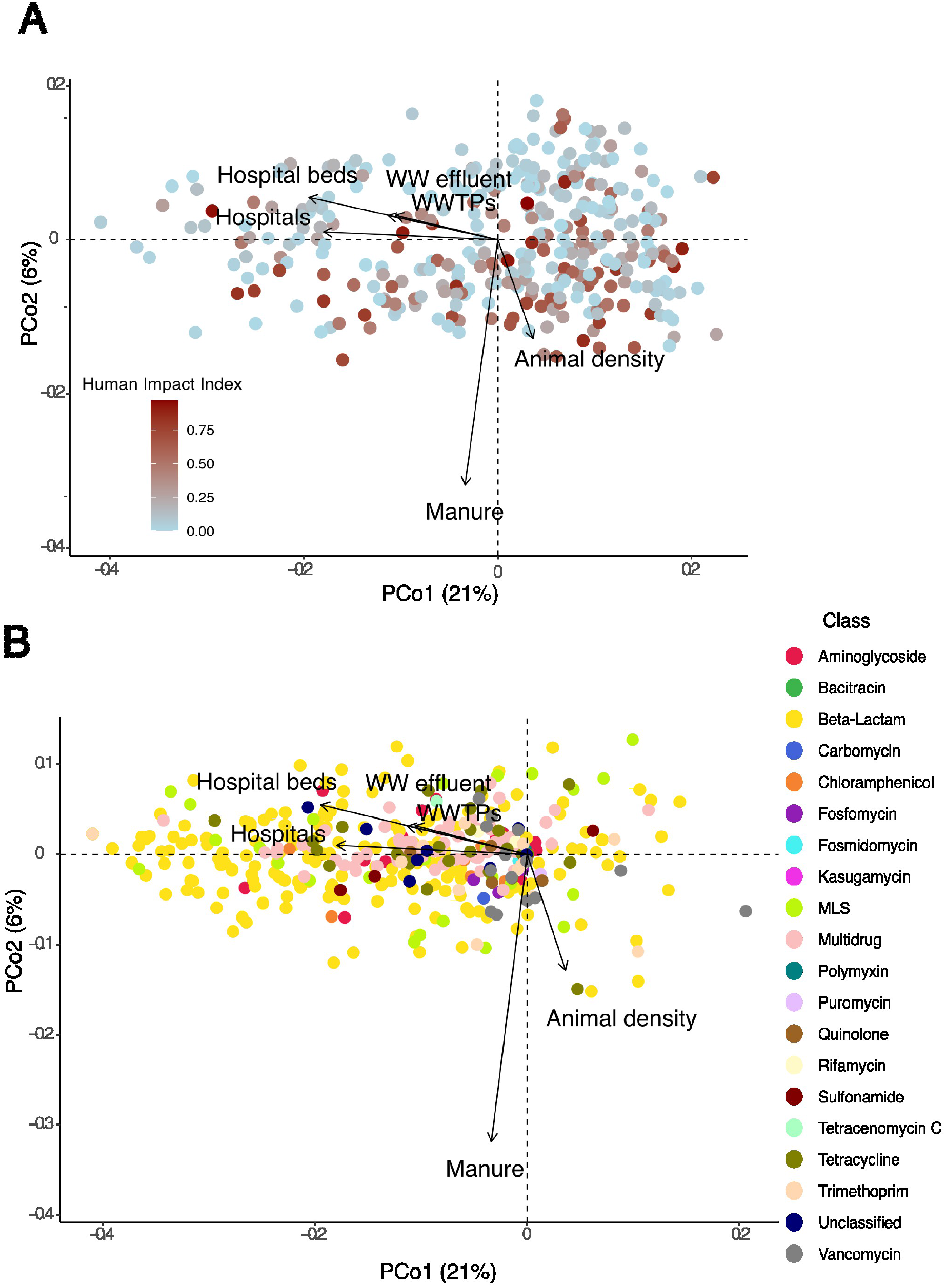
A) PCoA showing the distribution of 349 lake samples (based on presence-absence data) projected in the first two dimensions. Each sample is coloured by its HII. Arrows show variables that significantly correlated with the ordination after environmental fitting. B) PCoA showing the distribution of individual resistance genes (based on presence-absence data) projected in the first two dimensions. Each resistance gene is coloured by its class. Arrows show variables that significantly correlated with the ordination after environmental fitting.

### 3.4 Detection of antibiotic resistance genes in assembled data

Overall, we detected resistance gene-containing scaffolds in 246 out of the 349 lakes for a total of 1820 scaffolds. Of these, 661 scaffolds were longer than 1kb, and we assigned them to either plasmid or chromosomal origin. Most scaffolds containing resistance genes were assigned as chromosomal (598), whereas only 63 were characterized as plasmids. 98% (587) of chromosomal scaffolds and 84% (53) of plasmid-derived scaffolds could be taxonomically assigned to the genus-level using kraken2. The same taxa Mycolicibacterium (35% of chromosomal scaffolds / 17% of plasmid scaffolds), Acinetobacter (26% / 24%) and Mycobacterium (26% / 17%) contributed the most ARG-containing scaffolds on chromosomes or plasmids. Both Acinetobacter and Mycobacterium are widespread in the environment and sewage and have received attention as potential opportunistic pathogens (Doughari et al., 2009; Pfyffer, 2015). We subsequently tested whether the plasmid load (the count of ARG-containing scaffolds characterized as plasmid-originating divided by the total ARG-containing scaffolds in the sample) was significantly correlated with trophic state, watershed land use or resistance gene source variables, but found no indication (anova and Pearson’s product moment correlation tests, all *p* > 0.05).

### 3.5 Impact of land use and trophic status on the number of genomes

We tested whether trophic status or human impact index significantly correlated with the number of genomes detected in the samples. We did not detect a significant correlation between the average genome number estimated in a sample and the human impact index (Pearson’s product moment correlation test, *p* = 0.6), nor did we detect a significant impact of trophic status or human impact class (high, moderate, low) on the number of genomes detected (anova, all *p* > 0.05).

## 4. Discussion

Lakes integrate watershed-scale processes and are thus considered sentinels of processes such as climate change (Adrian et al., 2009), contaminant use (Schindler, 2009) and pathogen presence (Oliva et al., 2021). Lakes may thus constitute environmental receptors and reservoirs for ARGs, while at the same time providing critical ecosystem services, including 90% of Canadian drinking water (Environment Canada, 2000). Here, we take advantage of a continental-scale dataset of approximately 350 lakes to investigate whether we can detect signatures of human or veterinary use-induced antibiotic resistance across large spatial scales.

In contrast to previous work which used qPCR to detect absolute abundances of a small number of resistance genes (e.g., Czekalski et al., 2015; Pruden et al., 2012; Thevanon et al., 2012), we utilize metagenomic data, which is wider in scope, but produces relative abundances of ARGs. Thus, shifts in the total abundance of cells in a sample, for example during a bloom event under eutrophic conditions, could lead to a decrease in relative abundance of an ARG (if the blooming organisms are outcompeting the organisms containing the ARG), but could nonetheless represent a total increase of the ARG if measured via qPCR. We did not detect any indication that metagenomic data sets from lakes with higher human impact index or trophic status were characterized by a higher number of genomes, indicating that the relative abundances we report here may correlate well with absolute abundances of resistance genes.

Approximately 80 % of antibiotics sold in Canada are used in a veterinary context (Public Health Agency of Canada, 2022). The most commonly sold antibiotics for animals are tetracyclines, followed by macrolides, penicillins, sulfonamides and lincosamines (Public Health Agency of Canada, 2022). On the other hand, the most common ARGs detected in our dataset conferred resistance to bacitracin or were multidrug efflux pumps, followed by genes conferring resistance to macrolides, lincosamides and streptogramin. This partial mismatch between the antibiotics used and the genes conferring resistance to them points towards the complex roles that these genes play in microbial ecology. Moreover, links between antibiotics or antimicrobials and resistance genes might be further complicated by co-, and cross-selection (e.g., the use of ionophores, which are widely used in animals, but are not currently tracked as antibiotics (Wong, 2019).

Overall, the total relative abundance of ARGs was inversely correlated with total human watershed impact. This relationship was largely driven by the most abundant classes of genes conferring resistance against bacitracin and multidrug resistance genes. High relative abundances of genes conferring resistance to bacitracin have been previously identified in pristine lake sediments from a Tibetan lake (Chen et al., 2016), and in less-disturbed river samples (Lee et al., 2021). In contrast, Pan et al. (2020) detected an increase in the relative abundance of bacitracin ARGs with increasing lake nutrient status, but focused only on highly polluted lakes. Overall, the role of bacitracin resistance in aquatic bacterial ecology in the absence of direct anthropogenic pressures is unknown, but this resistance class may be unsuitable for environmental source tracking of ARG pollution (Lee et al., 2021).

The second most common class of resistance genes, multidrug resistance, is often associated with efflux pumps. Antibiotic resistance-related efflux pumps are prevalent both in antibiotic-resistant and sensitive bacteria, share homologies with other efflux pumps and resistance to an antibiotic may often be conferred via point mutations. Thus, multidrug efflux pumps may not be reliable markers of functional resistance (Nesme et al., 2014). We repeated the analyses after exclusion of all genes categorized with the mechanisms ‘efflux pumps’ or ‘membrane permeability change’, but found similar results, indicating that the misidentification of ARGs is not driving the patterns observed (Supplemental material).

The total relative abundance of resistance genes to other classes of antibiotics including aminoglycosides, fosmidomycin, puromycin, trimethoprim and vancomycin increased the higher human impact in the watershed. Trimethoprim, a synthetic antibiotic mostly used in human medicine, has been regularly detected in wastewater effluent and is present in lakes (Yang et al., 2018), where it is persistent and resistant to break-down (Bai and Acharya, 2017). However, the concentration in lakes is usually a magnitude lower (10-100 ng/L) than the MIC (100 µg/L), making *in situ* evolution of resistance unlikely (Yang et al., 2018). Therefore, ARGs likely enter the system via contamination with human wastewater. Genes conferring resistance to aminoglycosides have been previously found to be prevalent in interconnected lake and river systems (Zhang et al., 2021) and their prevalence in sediments has been previously linked to WWTP effluent (Devarajan et al., 2015). Vancomycin resistance is often associated with the use of veterinary antibiotics in animal production, from where it has spread to resistant pathogens such as vancomycin-resistant enterococci (VRE), which have been previously isolated from surface water (Messi et al., 2006). The genes detected in association with resistance to fosmidomycin and puromycin resistance all represent efflux pumps, and thus the same caveats may apply to them as to multidrug efflux pump resistance genes. However, they have been previously detected in stormwater (Białasek and Miłobędzka, 2020), and may thus be associated with urban impact on aquatic ecosystems. In summary, our findings support the notion that source tracking of anthropogenic pollution should be based on a multiple antibiotic resistance (MAR) framework, rather than individual ARGs (Kelsey et al., 2003; Krumperman, 1983).

The trophic state of a lake is often taken as an indicator of human activity. Even though we tried to reduce temporal variability by sampling all lakes at the height of summer stratification, lakes may be influenced by short term weather effects such as winds and precipitation. Taking into account the *in situ* trophic state of a lake at the time of sampling allows us to compare how well our GIS-data derived human impact index assessment matches the actual environment. We observed highly similar trends of resistance gene abundances whether we used human impact index or lake trophic state as an explanatory variable, indicating that GIS-derived land use data offers a valid proxy for *in situ* measurements to assess lake trophic states during summer stratification. Overall, resistance gene abundances were significantly different between trophic states, and significantly decreased from ultraoligotrophic to hypereutrophic lakes, due to the decrease of both bacitracin and multidrug resistance gene abundances with increasing trophic status. The overall relationship between lake nutrient status and the abundance of ARGs is unclear. Previous studies have reported positive correlations (Thevenon et al., 2012; Czekalski et al., 2015; Wang et al., 2020; Rajasekar et al., 2022) between select ARG absolute abundances and TP, but negative correlations have been reported as well (Huang et al., 2019). In contrast to previous results, we investigated the complete resistome (as represented by the CARD database) rather than a small selection of ARGs and extend the sampling beyond one or few lakes (which are often highly eutrophic) to encompass a broad variety of trophic statuses. Our results indicate that with increasing human activity and nutrient status the resistome shifts, with major ARG classes declining in relative abundance.

In correspondence with those shifts, we found that the overall composition of resistance genes to be significantly impacted by agricultural activity and pasture land use within the watershed, contrasting with the effects of natural landscapes and forestry. However, we did not detect an association of these significant land use variables with a specific class of resistance genes. Instead, the vast diversity of beta-lactam and multidrug resistance genes detected in the lake dataset was unassociated with any explanatory variable tested. Individual genes conferring resistance to trimethoprim, beta-lactams and MLS were associated with agricultural and pasture land use. This indicates that, while a large fraction of the lake resistome appears non-impacted by watershed land use, shifts from pristine or forestry-dominated (e.g., recent clear-cuts) watersheds to those associated with animal production do significantly alter resistome composition, likely in association with the veterinary use of antibiotics.

We specifically linked census-derived source variables regarding the origin of ARBs and ARGs to the composition of the resistome. Total population density, number of hospitals and hospital beds, the fraction of watershed with applied manure and the volume of wastewater effluent all significantly impacted ARG diversity, and the number of hospitals, hospital beds, and the volume of WWTP discharge were positively correlated with the total number of different ARGs detected. While this is an intriguing finding, we caution against over-interpretation, since only a small minority of lake watersheds contained any hospitals or WWTPs, and they often co-occur in the same watershed. Nonetheless, these correlations are in line with previous studies that found that WWTP effluent, and specifically effluent from hospitals, was enriched for resistance genes associated with clinical antibiotics usage (Reinthaler et al., 2003; Hocquet et al., 2016). Moreover, WWTP effluent has been identified as a point source for the pollution with ARGs and ARBs in previous work, including lakes in Switzerland (Czekalski et al., 2015) and lake Michigan (Chu et al., 2018), as well as river systems (Pei et al., 2006). The overall influence of WWTP discharge does not appear to be significant enough to cause shifts in overall ARG abundances in lakes but acts as a source for novel resistance genes in the system. Even though the overall impact of ARGs involved in human-associated antibiotics use is much smaller than that associated with animal production in the watershed, human-associated ARGs are of special concern for human health, due to a higher risk of reinfection, as well as the evolution of novel pathogens due to the dissemination of ARGs into new genetic backgrounds (Kraemer et al., 2019).

In general, human activity might alter the lake resistome directly, via the inflow of ARGs from wastewater or agricultural settings, or indirectly, via *in situ* evolution or community shifts caused by anthropogenic activity. Previous work has shown that environmental ARG abundances correlate well with markers of wastewater contamination in all except the most polluted environments, indicating the importance of direct effects on the resistome (Chen et al., 2019; Karkman et al., 2019).

While co-selection of antibiotic resistance with heavy metals has been widely reported (Yang et al., 2017; Rajasekar et al., 2022), we did not detect any indication of a correlation between mining within the watershed, and the putative contamination potential with heavy metals, and the resistome. This is likely because mining is very unevenly distributed across watersheds. While 26% of lakes have any watershed area influenced by mining, only 3% of lakes have more than 1% of their watershed impacted.

Previously, a highly impacted lake ecosystem has been shown to be enriched for mobile genetic elements, leading us to hypothesise that increasing human impact may select for an increased abundance of ARGs located on plasmids (Bengtsson-Palme et al., 2014). However, we did not detect any indication that land use, trophic status or resistance gene pollution sources altered the propensity of resistance genes to be located on chromosomes versus plasmids. This may be because the ecosystems studied are much less impacted than the polluted lake in the original study, both by direct pollution with antibiotics from antibiotics-producing industry, as well as with ARBs that may enter the system via wastewater. It is thus likely that the range of environmental conditions of the lakes described here is not extreme enough to exert the selective pressures necessary to shift towards a more mobile resistome (Bengtsson-Palme et al., 2017).

In summary, we present a survey of the resistome of 349 lakes ranging from pristine to highly impacted. To our knowledge, this is the largest systematic study of aquatic resistomes to date, which we aim to link to watershed land use, as well as sources of ARG point pollution. We detected a vast ‘natural’ aquatic resistome, which likely corresponds to ARGs used in inter-bacterial communication and competition. However, we also detected that the resistome was structured by watershed-scale land use, with pasture and agriculture-dominated lakes contrasting with those in pristine or forestry-dominated watersheds. Correspondingly, we did detect an impact of pollution sources for veterinary-origin ARGs on the structure of lake resistomes. Moreover, human antibiotics use within the watershed was found to have a positive correlation with ARG diversity. Taken together, these finding support the monitoring of ARG loading by both agricultural and clinical pollution sources in sink environments such as lakes.

