## Supplementary figures and images for "A resistome survey across hundreds of freshwater bacterial communities reveals the impacts of veterinary and human antibiotics use"

### Supplemental Figure 1

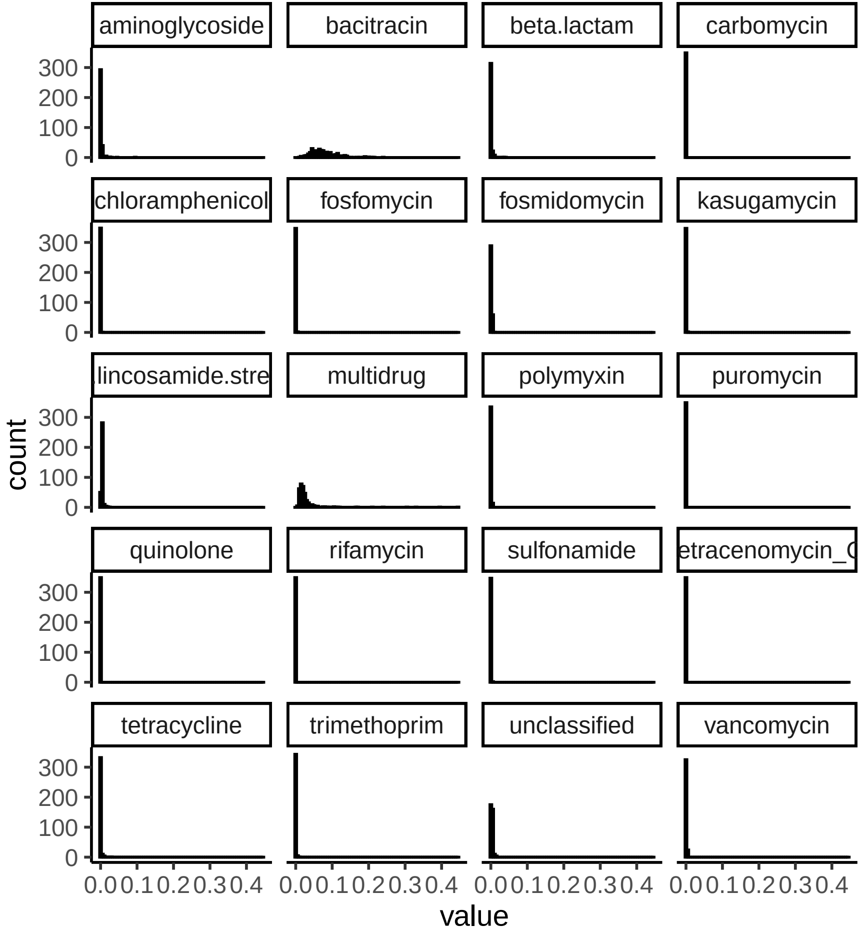
